## Supplemental Figs and Tables for "GATA6 regulates WNT and BMP programs to pattern precardiac mesoderm during the earliest stages of human cardiogenesis"

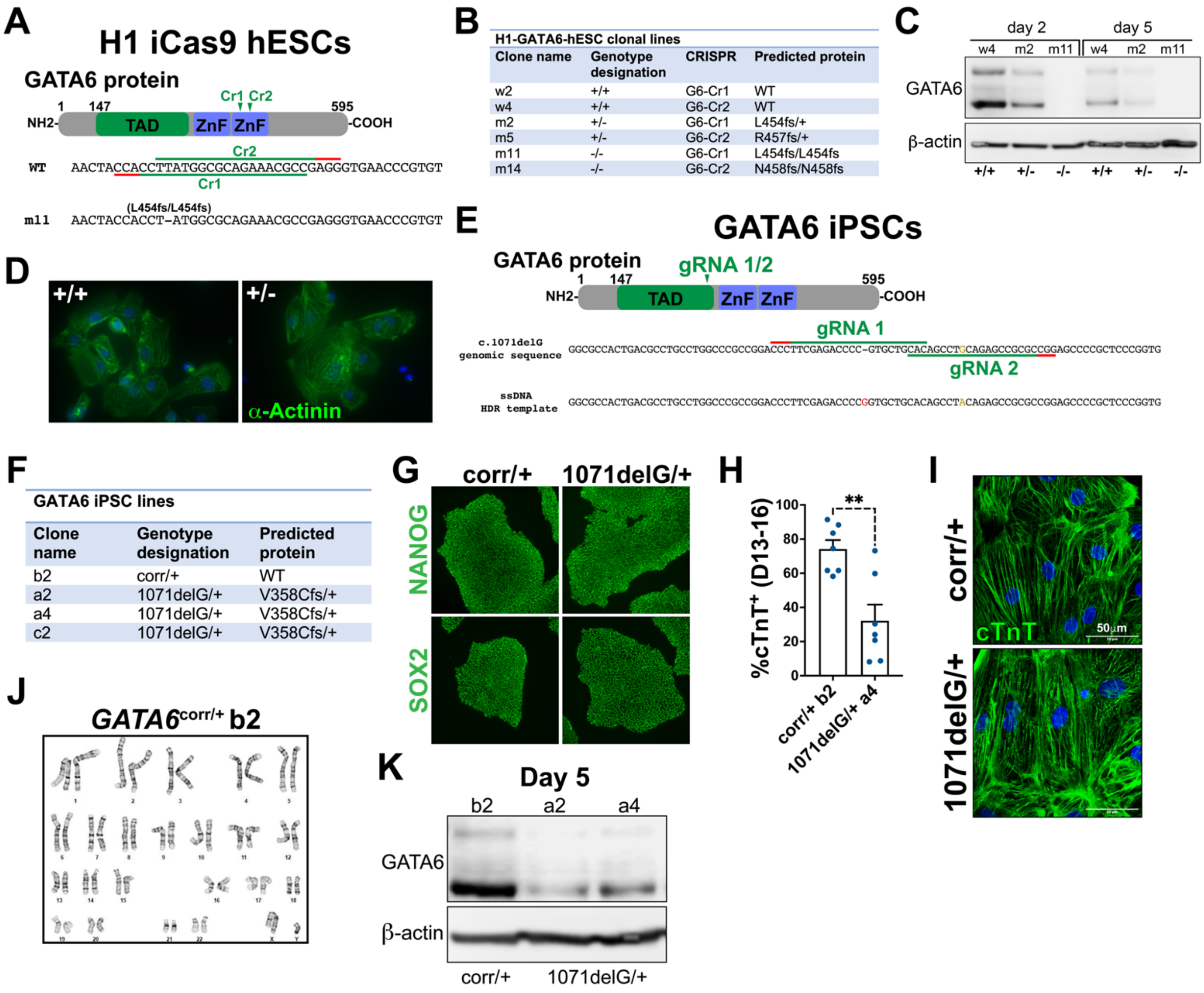


**H1-GATA6-hESC clonal lines**

| Clone name | Genotype designation | CRISPR | Predicted protein |
|---|---|---|---|
| w2 | +/+ | G6-Cr1 | WT |
| w4 | +/+ | G6-Cr2 | WT |
| m2 | +/- | G6-Cr1 | L454fs/+ |
| m5 | +/- | G6-Cr2 | R457fs/+ |
| m11 | -/- | G6-Cr1 | L454fs/L454fs |
| m14 | -/- | G6-Cr2 | N458fs/N458fs |

**GATA6 iPSC lines**

| Clone name | Genotype designation | Predicted protein |
|---|---|---|
| b2 | corr/+ | WT |
| a2 | 1071delG/+ | V358Cfs/+ |
| a4 | 1071delG/+ | V358Cfs/+ |
| c2 | 1071delG/+ | V358Cfs/+ |

**Supplemental Figure 1: CRISPR gene editing and characterization of mutant GATA6 hESC and iPSC lines.** (**A**) GATA6 CRISPR targeting scheme using H1 iCas9 hESCs was described in Shi et al. (Ref. 20). Two CRISPR gRNAs were used targeting the C-terminal zinc finger domain (GATA6-Cr1 and Cr2 gRNA target sequence in green, red indicates the PAM). (**B**) Table describing H1-GATA6-hESC clonal lines, genotype designation, and gRNA used in gene editing. Predicted protein describes mutant alleles according to the Human Genome Variation Society (HGVS) guidelines, fs indicates frameshift mutation. (**C**) Western blots from *GATA6*$^{+/+}$, *GATA6*$^{+/-}$, and *GATA6*$^{-/-}$ protein lysates at day 2 or 5 of cardiac differentiation probed for GATA6 with β-actin used as a loading control. (**D**) Immunofluorescence for the sarcomere marker α-Actinin and co-stained with DAPI on day 23 lactate-purified CMs. (**E**) Schematic for CRISPR-based correction of the GATA6 c.1071delG iPSC mutant allele to WT sequence (*GATA6*$^{corr/+}$). A pair of gRNAs flanking the G deletion site were used for targeting with Nickase-Cas9 and a WT repair template to allow for homology directed repair (HDR) of the mutant allele. A single base substitution of G to A (indicated by brown) was used in the WT repair template to induce a silent mutation in the gRNA 2 recognition sequence to prevent additional Cas9 activity after HDR. (**F**) Table describing the GATA6 iPSC clonal lines. The GATA6 c.1071delG mutant allele is indicated as the protein V358Cfs (according to HGVS guidelines). (**G**) Immunofluorescence for the pluripotency markers NANOG or SOX2 on

*GATA6*$^{corr/+}$ or *GATA6*$^{1071delG/+}$ iPSC colonies. (**H**) Flow cytometry quantification for %cTnT$^{+}$ CMs following cardiac directed differentiation for days 13 to 16 of *GATA6*$^{corr/+}$ or *GATA6*$^{1071delG/+}$ iPSCs. Data represents the mean ± SEM, dots represent independent biological replicates. Significance defined as **$p<0.01$ using the two-tailed Student's T-test. (**I**) Immunofluorescence for cTnT and co-stained with DAPI on day 23 lactate-purified iPSC-CMs. (**J**) Representative karyogram for *GATA6*$^{corr/+}$ iPSCs. (**K**) Western blots from *GATA6*$^{corr/+}$ or *GATA6*$^{1071delG/+}$ protein lysates at day 5 of cardiac differentiation probed for GATA6 with β-actin used as a loading control.

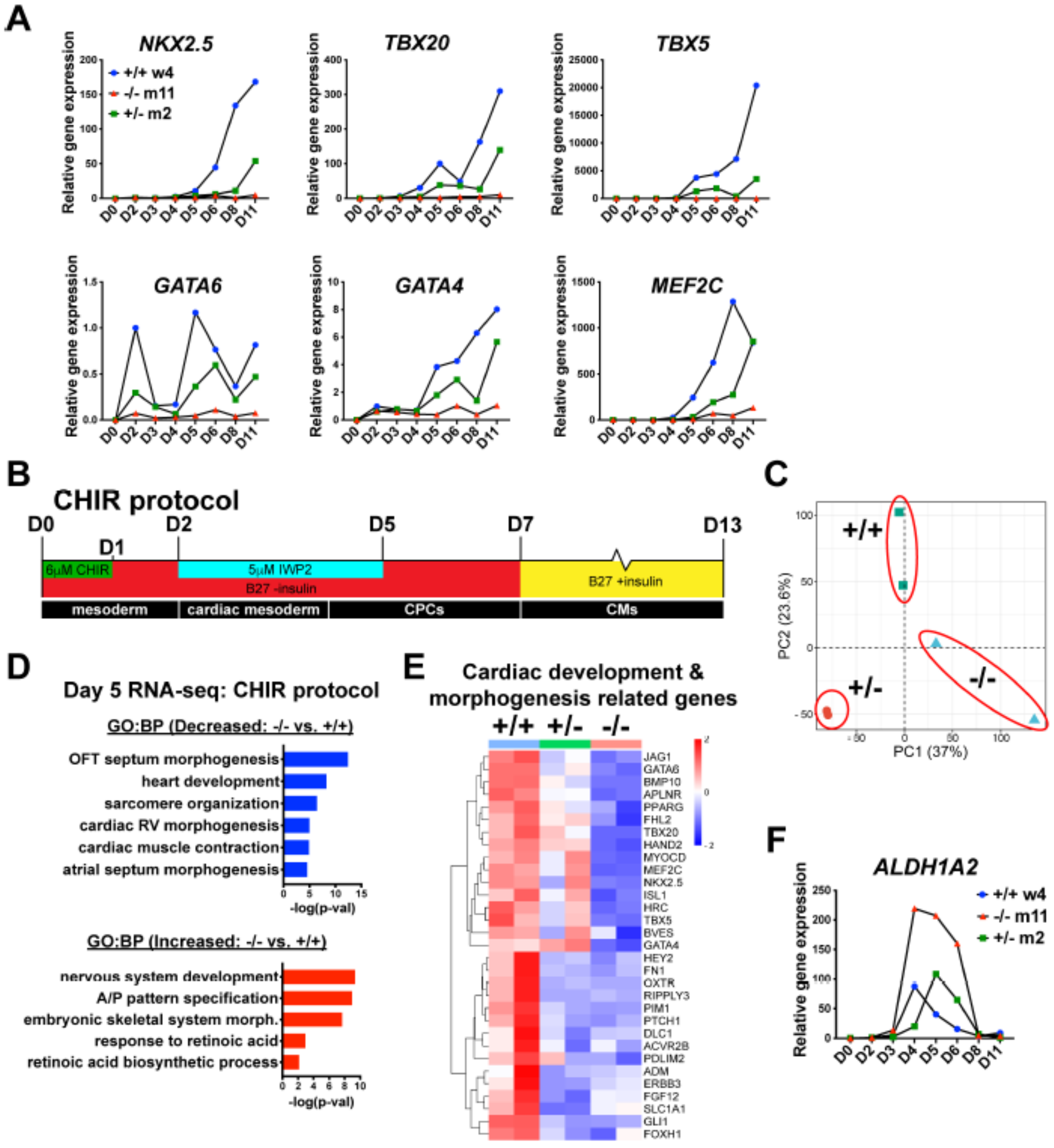


**Supplemental Figure 2: CPC marker gene transcriptional analysis of GATA6 WT or mutant hESCs.** (**A**) RT-qPCR time course for CPC markers (and *GATA6*) in differentiating *GATA6*$^{+/+}$, *GATA6*$^{+/-}$, and *GATA6*$^{-/-}$ hESCs (normalized to day 2 WT gene expression levels). (**B**) Schematic for *in vitro* CM directed differentiation CHIR protocol. (**C**) Principal component analysis (PCA) for comparisons of day 5 bulk RNA-seq from *GATA6*$^{+/+}$, *GATA6*$^{+/-}$, and *GATA6*$^{-/-}$ hESCs using the CHIR protocol (n=2). (**D**) Gene ontology (GO) analysis for biological process (BP) using differentially expressed genes (DEGs) from day 5 RNA-seq comparing *GATA6*$^{-/-}$ to WT. (**E**) Heatmap for cardiac development related genes from day 5 RNA-seq. Color gradient indicates relative gene expression level. (**F**) RT-qPCR time course for *ALDH1A2* transcript levels in differentiating *GATA6*$^{+/+}$, *GATA6*$^{+/-}$, and *GATA6*$^{-/-}$ hESCs using the cytokine-based protocol.

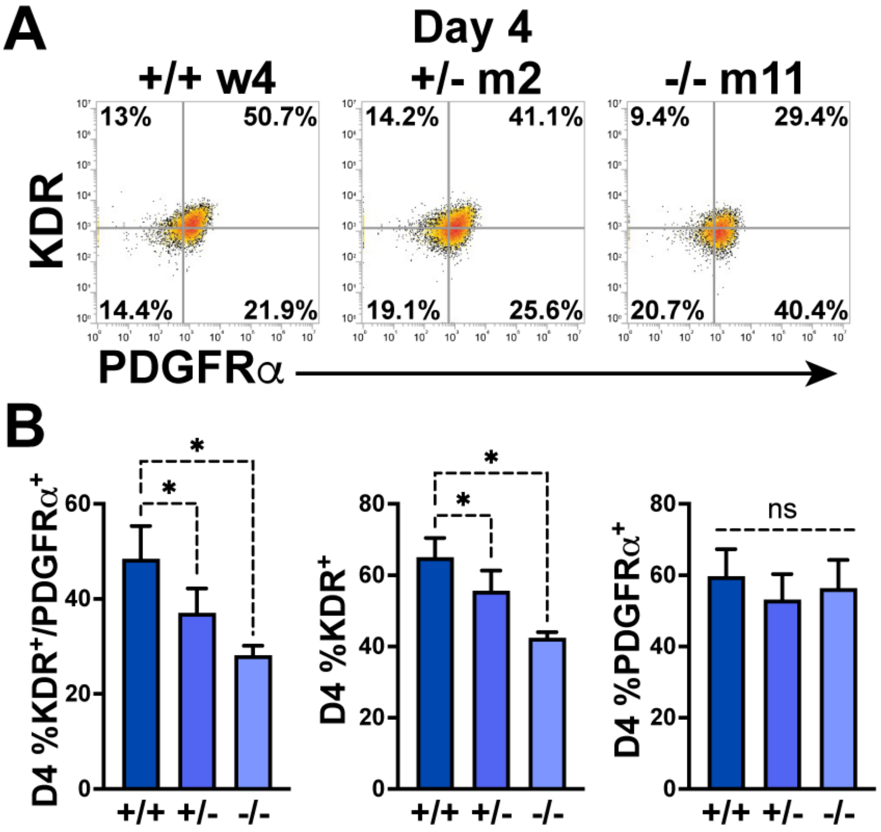


**Supplemental Figure 3: KDR and PDGFRα cardiac mesoderm analysis at day 4.** (**A**) Representative flow cytometry plots at day 4 of cardiac differentiation analyzing the % KDR and PDGFRα double-positive cells (%$K^+P^+$ ) from *GATA6*$^{+/+}$, *GATA6*$^{+/-}$, or *GATA6*$^{-/-}$ hESCs. (**B**) Quantification of day 4 flow cytometry data for %$K^+P^+$ double-positive, %$KDR^+$ single positive, or %PDGFR$\alpha^+$ single positive cells (n=6). Data represent the mean ± SEM, with statistical significance indicated as *$p<0.05$ and ns indicating not significant by one-way repeated measures ANOVA with Holm-Šídák's multiple comparison test.

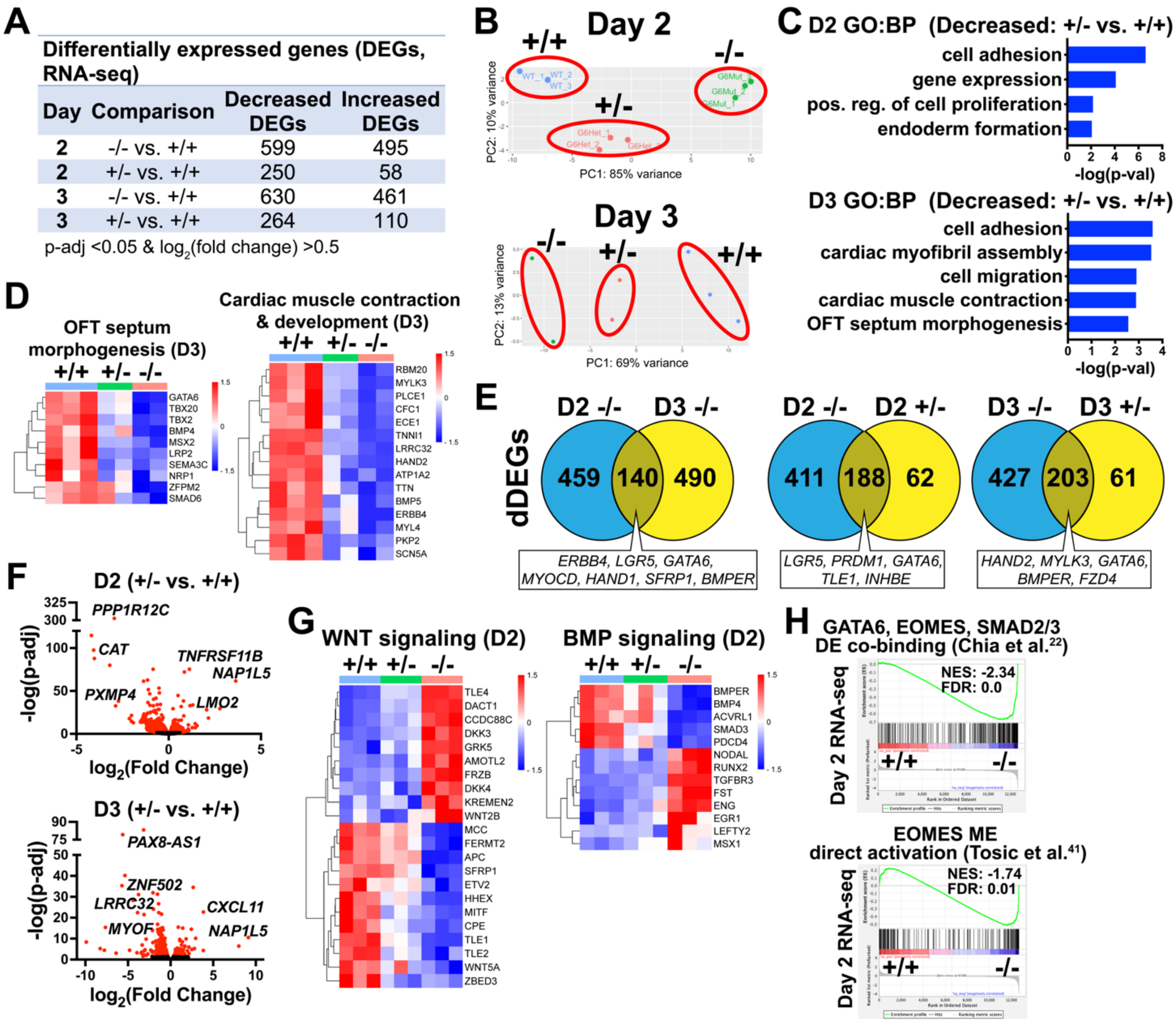


**Supplemental Figure 4: RNA-seq analysis following cardiac differentiation at day 2 or day 3.** (**A**) Number of DEGs identified when comparing *GATA6*$^{+/-}$ and *GATA6*$^{-/-}$ samples to *GATA6*$^{+/+}$ controls from day 2 and 3 RNA-seq data. **B**) PCA analysis for *GATA6*$^{+/+}$, *GATA6*$^{+/-}$, and *GATA6*$^{-/-}$ profiles from day 2 or day 3 RNA-seq. Data is representative of at least 2 independent biological replicates per sample. (**C**) Day 2 and 3 GO (BP) analysis using dDEGs identified comparing *GATA6*$^{+/-}$ to WT samples. (**D**) Heatmaps for OFT septum morphogenesis and cardiac development related genes from day 3 RNA-seq. Color gradient indicates relative gene expression levels. (**E**) Venn diagram comparisons of dDEGs (relative to WT) identified by day 2 or day 3 RNA-seq of *GATA6*$^{+/-}$ and *GATA6*$^{-/-}$ samples. (**F**) Volcano plots for day 2 or 3 *GATA6*$^{+/-}$ sample RNA-seq gene expression data relative to WT. Dots represent genes, red indicates p-adj<0.05 and black indicates p-adj>0.05. (**G**) Heatmaps for WNT and BMP related genes from day 2 RNA-seq. (**H**) Gene set enrichment analysis (GSEA) analysis of day 2 RNA-seq gene expression data (*GATA6*$^{-/-}$ relative to WT) using the GATA6, EOMES, and SMAD2/3 co-binding during hESC-DE differentiation dataset from Chia et al. (Ref. 22) and EOMES ME direct activation dataset from Tosic et al. (Ref. 41). NES indicates normalized enrichment score; FDR indicates false discovery rate.

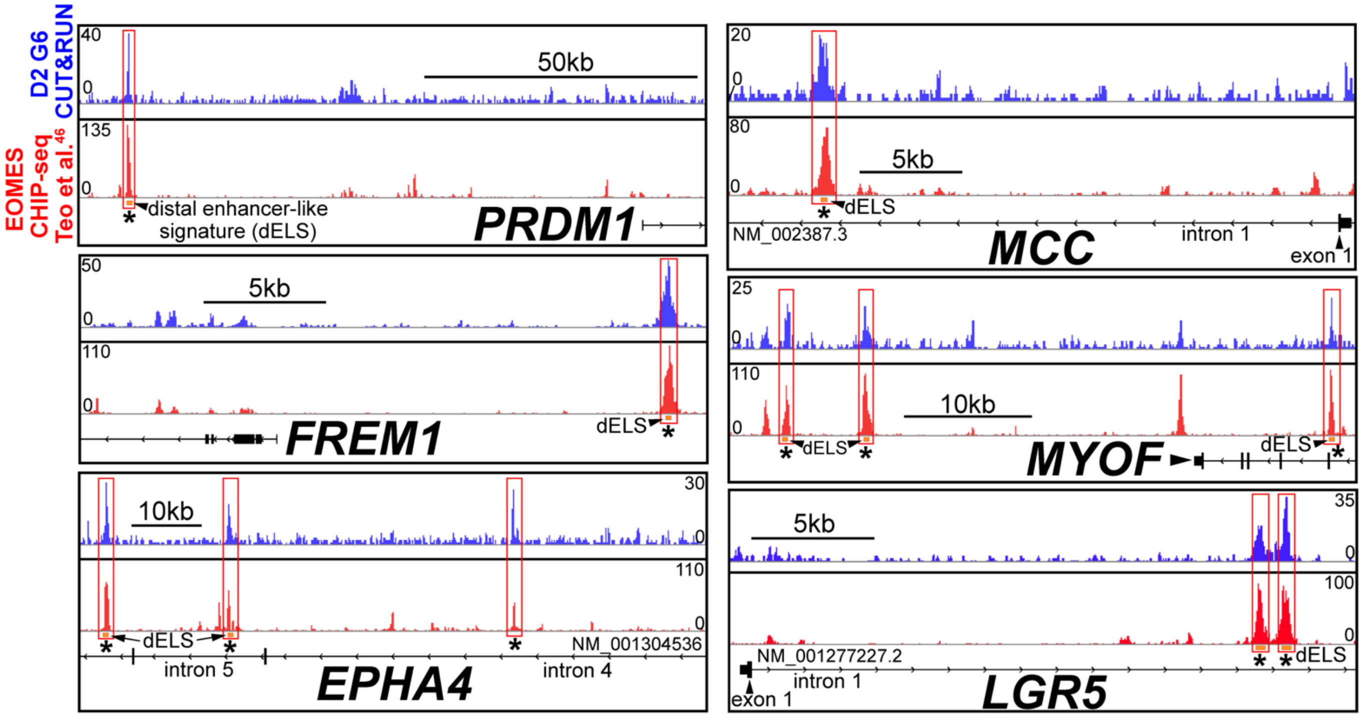


**Supplemental Figure 5: GATA6 CUT&RUN peaks overlap with previously published EOMES CHIP-seq peaks.** Human genome browser representations of GATA6 CUT&RUN data at day 2 of cardiac differentiation (blue tracks) and EOMES CHIP-seq identified genes at day 2 of hESC-DE differentiation from Teo et al. (Ref. 46) (red tracks). Orange rectangles represent approximate location for distal enhancer-like signatures (dELS) via the ENCODE Project (indicated by black arrows). Asterisks indicate significant GATA6 binding peaks ($p<0.003$ relative to IgG controls) and significant EOMES binding peaks as defined by Teo et al. (Ref. 46).

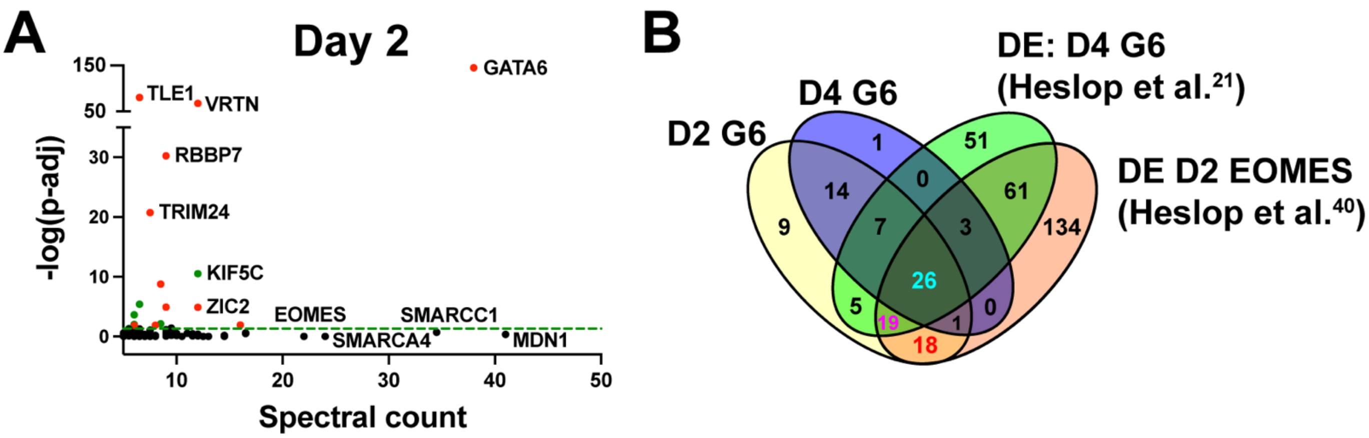


**C**

**Quadruple overlap**

| | | |
|---|---|---|
| GATA6 | HDAC2 | RBM39 |
| SMARCA4 | MTA1 | MTA2 |
| GTF2I | CBX1 | SMARCD1 |
| FUBP3 | CTBP2 | TRIM24 |
| MSH6 | DDX42 | HDAC1 |
| SSRP1 | GATAD2A | MBD3 |
| VRTN | MTA3 | MSH2 |
| KDM1A | RBBP7 | RBM4 |
| SALL4 | CTBP1 | |

**D2 G6 & DE: D2 EOMES[40]**

| | |
|---|---|
| ZIC2 | CECR2 |
| SMC3 | PTBP2 |
| SEPHS1 | PSMD5 |
| RCOR2 | RECQL |
| WDHD1 | SIN3A |
| MIXL1 | SMARCB1 |
| OGT | ARID1B |
| SMARCC2 | DNAJC8 |
| SART3 | RNF40 |

**D2 G6, DE: D4 G6[21], DE: D2 EOMES[40]**

| | |
|---|---|
| MDN1 | FEN1 |
| EOMES | SMARCE1 |
| SMC1A | TLE1 |
| NSUN2 | TRIM33 |
| XPO5 | U2SURP |
| ACTL6A | PSME3 |
| EP300 | SRRT |
| SMARCD2 | HAT1 |
| PPM1G | PRPF40A |
| BUB3 | |

**Supplemental Figure 6: Extended GATA6-RIME analysis.** (**A**) Graph depicts GATA6-RIME enriched proteins (spectral count) identified at day 2 of cardiac differentiation and their corresponding day 2 RNA-seq gene expression level significance (p-adj, *GATA6*$^{-/-}$ relative to WT). Dots indicate genes, black indicates p-adj>0.05, red indicates decreased expression level (p-adj<0.05), green indicates increased expression level (p-adj<0.5). (**B**) Venn diagram comparisons for GATA6-RIME enriched proteins from day 2 (D2 G6) and day 4 (D4 G6) of cardiac differentiation from the present study with day 4 GATA6-RIME (DE: D4 G6) or day 2 EOMES-RIME (DE: D2 EOMES) enriched proteins reported during hESC-DE differentiation from Heslop et al. (Ref. 21 and Ref. 39). (**C**) Proteins identified in the Venn diagram comparisons of RIME data described in (B).

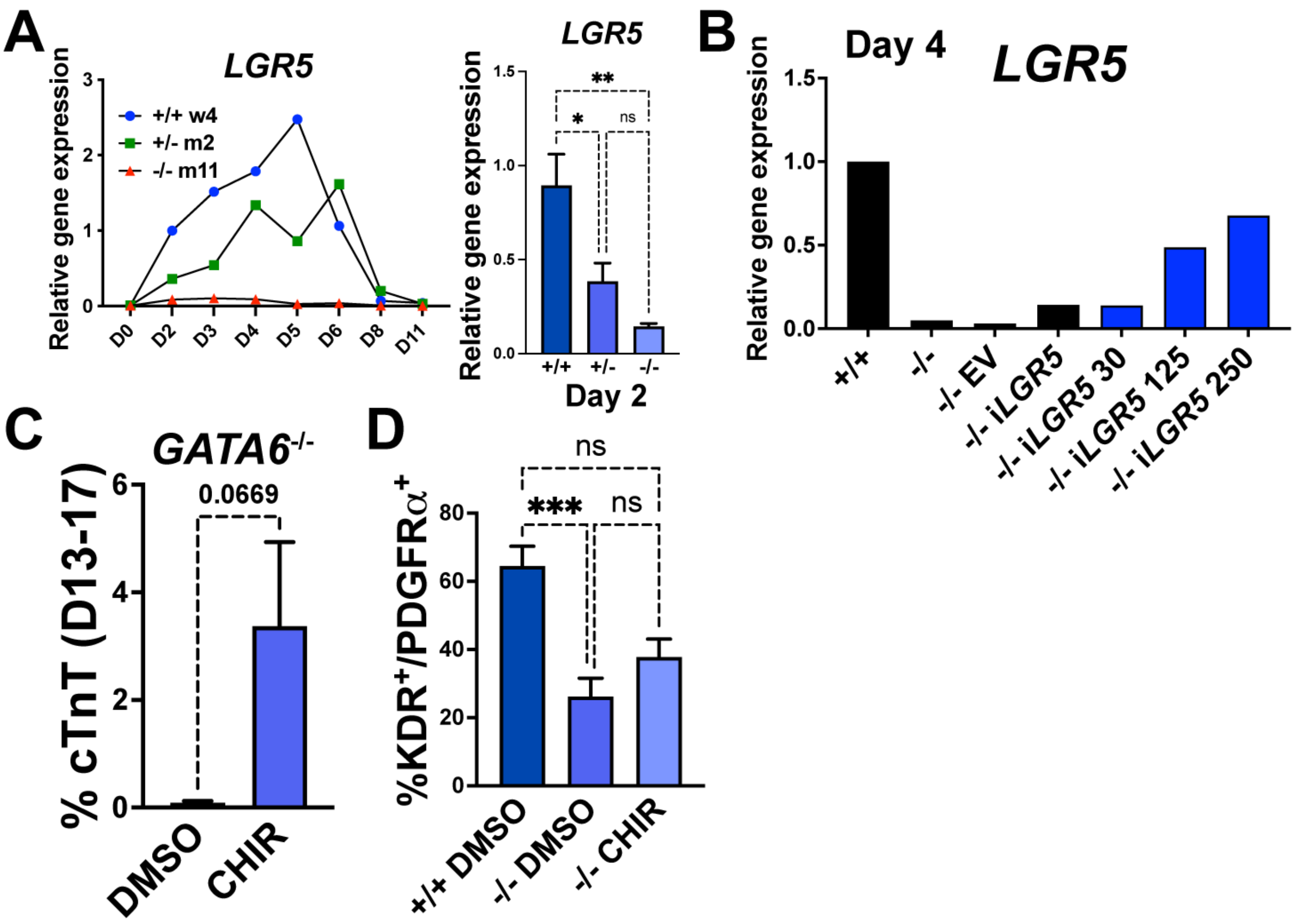


**Supplemental Figure 7: Extended data for i*LGR5* and early CHIR treatment.** (**A**) RT-qPCR time course for relative *LGR5* expression in differentiating *GATA6*$^{+/+}$, *GATA6*$^{+/-}$, and *GATA6*$^{-/-}$ hESCs normalized to day 2 WT gene expression level (on left). Day 2 RT-qPCR quantification for relative *LGR5* expression level normalized to WT (on right, n=6). (**B**) Day 4 RT-qPCR for relative *LGR5* expression level (normalized to WT) in i*LGR5* or EV transduced *GATA6*$^{-/-}$ hESCs treated with or without varying concentrations of DOX (30, 125, and 250 indicate ng/mL concentrations of DOX). (**C**) %cTnT$^{+}$ CMs from days 13 to 17 of cardiac differentiation quantified by flow cytometry in *GATA6*$^{-/-}$ cells treated with CHIR (3μM) or vehicle (n≥8). (**D**) Flow cytometry quantification for %K$^{+}$P$^{+}$ double-positive cells at day 5 of cardiac differentiation from *GATA6*$^{-/-}$ hESCs treated with CHIR and vehicle treated *GATA6*$^{+/+}$ or *GATA6*$^{-/-}$ hESCs controls (n≥3). Data represents the mean ± SEM, significance indicated by *p<0.05, **p<0.01, ***p<0.001, by two-tailed Student's t-test (C) or one-way ANOVA with Tukey's multiple comparisons test (A, D).

### Supplemental Table 1: GATA6-RIME Enriched Proteins

| 99 common elements in Day_2_G6_R1 and Day_2_G6_R2 | | |
|---|---|---|
| **Gene Name** | **Protein Name** | **Spectral Count (Avg)** |
| MDN1 | Midasin | 41 |
| GATA6 | Transcription factor GATA-6 | 38 |
| SMARCC1 | SWI/SNF complex subunit SMARCC1 | 34.5 |
| SMARCA4 | Transcription activator BRG1 | 24 |
| EOMES | Eomesodermin homolog | 22 |
| KLC3 | Kinesin light chain 3 | 16.5 |
| MRPL15 | 39S ribosomal protein L15 mitochondrial | 16.5 |
| DZIP3 | E3 ubiquitin-protein ligase DZIP3 | 16 |
| DNM2 | Dynamin-2 | 14.5 |
| GTF2I | General transcription factor II-I | 14.5 |
| FUBP3 | Far upstream element-binding protein 3 | 13 |
| KIF5B | Kinesin-1 heavy chain | 13 |
| MSH6 | DNA mismatch repair protein Msh6 | 12.5 |
| SSRP1 | FACT complex subunit SSRP1 | 12.5 |
| ARID1A | AT-rich interactive domain-containing protein 1A | 12 |
| GADD45GIP1 | Growth arrest and DNA damage-inducible proteins-interacting protein 1 | 12 |
| KIF5C | Kinesin heavy chain isoform 5C | 12 |
| VRTN | Vertnin | 12 |
| ZIC2 | Zinc finger protein ZIC 2 | 12 |
| KDM1A | Lysine-specific histone demethylase 1A | 11.5 |
| SALL4 | Sal-like protein 4 | 11.5 |
| TOP2B | DNA topoisomerase 2 (Fragment) | 11.5 |
| HDAC2 | Histone deacetylase 2 | 11 |
| MRPL44 | 39S ribosomal protein L44 mitochondrial | 10.5 |
| ELAVL1 | ELAV-like protein 1 | 10 |
| MTA1 | Metastasis-associated protein MTA1 | 10 |
| SMC1A | Structural maintenance of chromosomes protein | 10 |
| CBX1 | Chromobox protein homolog 1 | 9.5 |
| PYCR2 | Pyrroline-5-carboxylate reductase 2 | 9.5 |
| SMARCA5 | SWI/SNF-related matrix-associated actin-dependent regulator of chromatin subfamily A member 5 | 9.5 |
| SMC3 | Structural maintenance of chromosomes protein 3 | 9.5 |
| CTBP2 | C-terminal-binding protein 2 | 9 |
| DDX42 | ATP-dependent RNA helicase DDX42 | 9 |
| GATAD2A | Transcriptional repressor p66-alpha | 9 |
| MTA3 | Metastasis-associated protein MTA3 | 9 |
| NSUN2 | tRNA (cytosine(34)-C(5))-methyltransferase | 9 |
| RBBP7 | Histone-binding protein RBBP7 | 9 |
| SF1 | Splicing factor 1 | 9 |
| SMU1 | WD40 repeat-containing protein SMU1 | 9 |
| XPO5 | Exportin-5 | 9 |
| ACTL6A | Actin-like protein 6A | 8.5 |
| SEPHS1 | Selenide water dikinase 1 | 8.5 |
| CTBP1 | C-terminal-binding protein 1 | 8 |
| EP300 | Histone acetyltransferase p300 | 8 |
| RBM39 | RNA-binding protein 39 | 8 |
| RCOR2 | REST corepressor 2 | 8 |
| SMARCD2 | SMARCD2 protein | 8 |
| DAZAP1 | DAZ-associated protein 1 | 7.5 |
| HNRNPUL1 | Heterogeneous nuclear ribonucleoprotein U-like protein 1 | 7.5 |
| MRPL4 | 39S ribosomal protein L4 mitochondrial (Fragment) | 7.5 |
| MTA2 | Metastasis-associated protein MTA2 | 7.5 |
| SMARCD1 | SWI/SNF-related matrix-associated actin-dependent regulator of chromatin subfamily D member 1 | 7.5 |
| TRIM24 | Transcription intermediary factor 1-alpha | 7.5 |
| WDHD1 | WD repeat and HMG-box DNA-binding protein 1 | 7.5 |
| HDAC1 | Histone deacetylase 1 | 7 |
| MIXL1 | Homeobox protein MIXL1 | 7 |
| OGT | UDP-N-acetylglucosamine--peptide N-acetylglucosaminyltransferase 110 kDa subunit | 7 |
| PPM1G | Protein phosphatase 1G | 7 |
| SMARCC2 | SWI/SNF complex subunit SMARCC2 | 7 |
| XRN2 | 5'-3' exoribonuclease 2 | 7 |
| BUB3 | Mitotic checkpoint protein BUB3 (Fragment) | 6.5 |
| DUT | DUTP pyrophosphatase isoform CRA_c | 6.5 |
| FEN1 | Flap endonuclease 1 | 6.5 |
| MBD3 | Methyl-CpG-binding domain protein 3 | 6.5 |
| MSH2 | DNA mismatch repair protein Msh2 | 6.5 |
| RBM4 | RNA-binding protein 4 | 6.5 |
| SART3 | Squamous cell carcinoma antigen recognized by T-cells 3 | 6.5 |
| SMARCE1 | SWI/SNF-related matrix-associated actin-dependent regulator of chromatin subfamily E member 1 (Fragment) | 6.5 |
| SNX9 | Sorting nexin-9 | 6.5 |
| THRAP3 | Thyroid hormone receptor-associated protein 3 | 6.5 |
| TLE1 | Transducin-like enhancer protein 1 | 6.5 |
| TRIM33 | E3 ubiquitin-protein ligase TRIM33 (Fragment) | 6.5 |
| U2SURP | U2 snRNP-associated SURP motif-containing protein | 6.5 |
| CECR2 | Cat eye syndrome critical region protein 2 | 6 |
| MRPL38 | 39S ribosomal protein L38 mitochondrial | 6 |
| MRPL47 | 39S ribosomal protein L47 mitochondrial | 6 |
| PSME3 | Proteasome activator complex subunit 3 | 6 |
| PTBP2 | Polypyrimidine tract-binding protein 2 | 6 |
| SRRT | Serrate RNA effector molecule homolog | 6 |
| USP9X | Probable ubiquitin carboxyl-terminal hydrolase FAF-X | 6 |
| DDX46 | Probable ATP-dependent RNA helicase DDX46 | 5.5 |
| HAT1 | Histone acetyltransferase type B catalytic subunit | 5.5 |
| HINT1 | Histidine triad nucleotide-binding protein 1 | 5.5 |
| MRPL13 | 39S ribosomal protein L13 mitochondrial | 5.5 |
| MRPL20 | 39S ribosomal protein L20 mitochondrial | 5.5 |
| MRPL50 | 39S ribosomal protein L50 mitochondrial | 5.5 |
| MSI1 | RNA-binding protein Musashi homolog 1 | 5.5 |
| PSMD5 | 26S proteasome non-ATPase regulatory subunit 5 | 5.5 |
| RECQL | ATP-dependent DNA helicase Q1 | 5.5 |
| SIN3A | Paired amphipathic helix protein Sin3a | 5.5 |
| SMARCB1 | SWI/SNF related matrix associated actin dependent regulator of chromatin subfamily b member 1 isoform CRA_c | 5.5 |
| SNRNP70 | U1 small nuclear ribonucleoprotein 70 kDa | 5.5 |
| ARID1B | AT-rich interactive domain-containing protein 1B | 5 |
| DNAJC8 | DnaJ homolog subfamily C member 8 | 5 |
| MRPL43 | 39S ribosomal protein L43 mitochondrial (Fragment) | 5 |
| MRPL49 | 39S ribosomal protein L49 mitochondrial | 5 |
| MSI2 | RNA-binding protein Musashi homolog 2 | 5 |
| PRPF40A | Pre-mRNA-processing factor 40 homolog A | 5 |
| RNF40 | E3 ubiquitin-protein ligase BRE1B | 5 |
| RNF40 | E3 ubiquitin-protein ligase BRE1B | 5 |

| 52 common elements in Day_4_G6_R1 and Day_4_G6_R2 | | |
|---|---|---|
| **Gene Name** | **Protein Name** | **Spectral Count (Avg)** |
| DNM2 | Dynamin-2 | 24.5 |
| GATA6 | Transcription factor GATA-6 | 24 |
| SMARCC1 | SWI/SNF complex subunit SMARCC1 | 24 |
| PYCR2 | Pyrroline-5-carboxylate reductase 2 | 23.5 |
| DZIP3 | E3 ubiquitin-protein ligase DZIP3 | 21.5 |
| SMARCA4 | Transcription activator BRG1 | 18.5 |
| FUBP3 | Far upstream element-binding protein 3 | 15.5 |
| MRPL15 | 39S ribosomal protein L15 mitochondrial | 13 |
| MRPL44 | 39S ribosomal protein L44 mitochondrial | 13 |
| SNX9 | Sorting nexin-9 | 12 |
| GADD45GIP1 | Growth arrest and DNA damage-inducible proteins-interacting protein 1 | 11.5 |
| SSRP1 | FACT complex subunit SSRP1 | 11.5 |
| KIF5B | Kinesin-1 heavy chain | 10.5 |
| MSH6 | DNA mismatch repair protein Msh6 | 9.5 |
| SALL4 | Sal-like protein 4 | 9.5 |
| VRTN | Vertnin | 9.5 |
| GTF2I | General transcription factor II-I | 9 |
| TRIM24 | Transcription intermediary factor 1-alpha | 9 |
| ELAVL1 | ELAV-like protein 1 | 8.5 |
| KDM1A | Lysine-specific histone demethylase 1A | 8.5 |
| SMU1 | WD40 repeat-containing protein SMU1 | 8.5 |
| HDAC2 | Histone deacetylase 2 | 8 |
| MSI2 | RNA-binding protein Musashi homolog 2 | 7.5 |
| MTA1 | Metastasis-associated protein MTA1 | 7.5 |
| RBBP7 | Histone-binding protein RBBP7 | 7.5 |
| SMARCD1 | SWI/SNF-related matrix-associated actin-dependent regulator of chromatin subfamily D member 1 | 7.5 |
| CBX1 | Chromobox protein homolog 1 | 7 |
| CTBP1 | C-terminal-binding protein 1 | 7 |
| DDX42 | ATP-dependent RNA helicase DDX42 | 7 |
| KLC3 | Kinesin light chain 3 | 7 |
| MSH2 | DNA mismatch repair protein Msh2 | 7 |
| MSI1 | RNA-binding protein Musashi homolog 1 | 7 |
| MTA3 | Metastasis-associated protein MTA3 | 7 |
| SF1 | Splicing factor 1 | 7 |
| TAF15 | TATA-binding protein-associated factor 2N | 7 |
| CTBP2 | C-terminal-binding protein 2 | 6.5 |
| RBM39 | RNA-binding protein 39 | 6.5 |
| SNRPN | Small nuclear ribonucleoprotein-associated protein N (Fragment) | 6.5 |
| TIAL1 | Nucleolysin TIAR | 6.5 |
| GATAD2A | Transcriptional repressor p66-alpha | 6 |
| MRPL47 | 39S ribosomal protein L47 mitochondrial | 6 |
| MTA2 | Metastasis-associated protein MTA2 | 6 |
| RBM4 | RNA-binding protein 4 | 6 |
| TOP2B | DNA topoisomerase 2 (Fragment) | 6 |
| USP9X | Probable ubiquitin carboxyl-terminal hydrolase FAF-X | 6 |
| ARID1A | AT-rich interactive domain-containing protein 1A | 5.5 |
| HDAC1 | Histone deacetylase 1 | 5.5 |
| HNRNPUL1 | Heterogeneous nuclear ribonucleoprotein U-like protein 1 | 5.5 |
| MRPL4 | 39S ribosomal protein L4 mitochondrial (Fragment) | 5.5 |
| ADAR | Double-stranded RNA-specific adenosine deaminase | 5 |
| MBD3 | Methyl-CpG-binding domain protein 3 | 5 |
| MRPL49 | 39S ribosomal protein L49 mitochondrial | 5 |

#### Supplemental Table 2: Cell lines, primer sequences, and antibodies used.

##### Cell Lines

| Name | Source | Notes |
|---|---|---|
| H1-GATA6-hESCs | Parent line: WiCell Research Institute, H1 (WA01), NIHhESC-10-0043 | Male |
| GATA6 c.1071delG iPSCs | Yu et al., 2014 (Ref: 15) | Male |
| 293T | ATCC: human embryonic kidney cells, CRL-11268 | |
| CF1 Mouse Embryonic Fibroblasts, irradiated | MTI Global Stem: CF-1 MEF 2M IRR | |

##### gRNAs sequences

| Name | Sequence 5' to 3' | Cell line used | Notes |
|---|---|---|---|
| G6-Cr1 | GGCGTTTCTGCGCCATAAGG | H1-GATA6-hESCs | |
| G6-Cr2 | TTATGGCGCAGAAACGCCG | H1-GATA6-hESCs | |
| gRNA 1 - F | caccgCACAGCCTGCAGAGCCGCGC | GATA6 c.1071delG iPSCs | gRNA target sequence capitalized |
| gRNA 1 - R | aaacGCGCGGCTCTGCAGGCTGTGc | GATA6 c.1071delG iPSCs | gRNA target sequence capitalized |
| gRNA 2 - F | caccgGTGCAGCACGGGGTCTCGAA | GATA6 c.1071delG iPSCs | gRNA target sequence capitalized |
| gRNA 2 - R | aaacTTCGAGACCCCGTGCTGCAc | GATA6 c.1071delG iPSCs | gRNA target sequence capitalized |

##### Primer sequences

| Name | Sequence 5' to 3' | Application |
|---|---|---|
| G6-iPSC-F | GCCCCTACTCGCCCTACG | PCR genotyping |
| G6-iPSC-R | CACCCGGACGAGAAAGTCCT | PCR genotyping |
| LGR5-cloning-F | TTCGTACGTACTTCGGGCACCATGGA | Full Length LGR5 PCR (using day 2 cDNA). Includes BsiWI site. |
| LGR5-cloning-R | GGCAATTGTTAGAGACATGGGACAAATGCC | Full Length LGR5 PCR (using day 2 cDNA). Includes MfeI site. |
| LGR5-seq1 | GAAAGAAATGCTTTGATGGG | LGR5 Sanger sequencing primer 1 |
| LGR5-seq2 | ATGCATTTTCCACTTTGCCA | LGR5 Sanger sequencing primer 2 |
| LGR5-seq3 | CTGGTGGGAGAATGGGGTTG | LGR5 Sanger sequencing primer 3 |
| CS-TRE-F | ATTTTCGGGTTTATTACAGG | Lentiviral vector Sanger sequencing primer |
| CS-TRE-R | GGAAAGGAGCTGACAGGT | Lentiviral vector Sanger sequencing primer |
| HPRT-F | ACCAGTCAACAGGGGACATAA | RT-qPCR |
| HPRT-R | CTTCGTGGGGTCCTTTTCACC | RT-qPCR |
| NKX2.5-F | AGCCGAAAAGAAAGAGCTGTGCG | RT-qPCR |
| NKX2.5-R | GACCTGCGCCTGCGAGAAGAG | RT-qPCR |
| TBX20-F | GGCGACGGAGAACACAATCAA | RT-qPCR |
| TBX20-R | CTGGGCACAGGACGACTTC | RT-qPCR |
| TBX5-F | AGCAGTGACTTCCTACCAGAAC | RT-qPCR |
| TBX5-R | TGACATTCTGTGCAGCTCCAT | RT-qPCR |
| GATA6-F | CTGCGGGCTCTACAGCAAG | RT-qPCR |
| GATA6-R | GTTGGCACAGGACAATCCAAG | RT-qPCR |
| GATA4-F | AAAGAGGGGATCCAAACCAG | RT-qPCR |
| GATA4-R | TTGCTGGAGTTGCTGGAAG | RT-qPCR |
| MEF2C-F | CTGGTGTAACACATCGACCTC | RT-qPCR |
| MEF2C-R | GATTGCCATACCCGTTCCCT | RT-qPCR |
| ALDH1A2-F | GGAGTCCCTTTGACCCCACCACT | RT-qPCR |
| ALDH1A2-R | CCCTTTCGGCCCAGTCCTTTGC | RT-qPCR |
| LGR5-F | GTTTCCCGCAAGACGTAACT | RT-qPCR |
| LGR5-R | CAGCGTCTTCACCTCCTACC | RT-qPCR |
| T-F | ACCCAGTTCATAGCGGTGAC | RT-qPCR |
| T-R | CCATTGGGAGTACCCAGGTT | RT-qPCR |
| EOMES-F | CTGCCCACTACAATGTGTTCG | RT-qPCR |
| EOMES-R | GCGCCTTTGTTATTGGTGAGTTT | RT-qPCR |

##### Antibodies

| Antibody | Source | Application |
|---|---|---|
| mouse monoclonal α-Actinin | Sigma-Aldrich A7811 | immunocytochemistry 1:200 |
| rabbit polyclonal NANOG | Cell Signaling Technology 3850 | immunocytochemistry 1:250 |
| rabbit polyclonal SOX2 | Invitrogen 48-1400 | immunocytochemistry 1:100 |
| mouse monoclonal cardiac troponin T | Invitrogen MA5-12960 | immunocytochemistry 1:200, flow cytometry 1:500 |
| rabbit monoclonal GATA6 | Cell Signaling Technology 5851 | western blotting 1:1000 |
| mouse monoclonal β-actin | Sigma-Aldrich A1978 | western blotting 1:50000 |
| mouse monoclonal anti-human KDR PE conjugated | R&D Systems FAB357P-100 | flow cytometry 10μL per million cells |
| mouse monoclonal anti-human PDGFRα PE conjugated | R&D Systems FAB1264A | flow cytometry 10μL per million cells |
| goat polyclonal anti-human/mouse Brachyury PE-conjugated | R&D Systems IC2085P | flow cytometry 1:40 |
| rabbit polyclonal SMAD2/3 | Cell Signaling Technology 3102 | western blotting 1:500 |
| rabbit monoclonal Phospho-SMAD2 (Ser465/467)/SMAD3 (Ser423/425) (D27F4) | Cell Signaling Technology 8828 | western blotting 1:500 |
| rabbit monoclonal Phospho-SMAD1 (Ser463/465)/ SMAD5 (Ser463/465)/ SMAD9 (Ser465/467) (D5B10) | Cell Signaling Technology 13820 | western blotting 1:500 |
| rabbit monoclonal non-phospho (Active) β-Catenin (Ser33/37/Thr41) (D13A1) | Cell Signaling Technology 8814 | western blotting 1:1000 |
| rabbit polyclonal β-Catenin | Cell Signaling Technology 9562 | western blotting 1:1000 |
| rabbit monoclonal SMARCC1/BAF155 (D7F8S) | Cell Signaling Technology 11956 | western blotting 1:1000 |
| rabbit polyclonal TBR2 / Eomes | Abcam ab23345 | western blotting 1:500 |
| goat anti-mouse IgG1 Alexa Fluor™ 488 | Invitrogen A-21121 | immunocytochemistry 1:500 |
| goat anti-rabbit Alexa Fluor™ 488 | Invitrogen A-11008 | immunocytochemistry 1:1000 |
| goat anti-mouse Alexa Fluor™ 488 | Invitrogen A-10667 | immunocytochemistry 1:500 |
| goat anti-mouse Alexa Fluor™ 647 | Invitrogen A-21236 | flow cytometry 1:400 |
| goat anti-rabbit HRP | Bio-Rad 1706515 | western blotting 1:2000 - 1:10000 |
| goat anti-mouse HRP | Bio-Rad 1706516 | western blotting 1:2000 - 1:10000 |
| mouse monoclonal IgG1 APC-conjugated | R&D Systems IC002A | flow cytometry 10μL per million cells |
| mouse monoclonal IgG1 PE-conjugated | R&D Systems IC002P | flow cytometry 10μL per million cells |
| goat polyclonal IgG PE-conjugated | R&D Systems IC108P | flow cytometry 1:40 |
